# Specific dendritic cells spatial organization is associated to ICB Response in Non–Small-Cell Lung Cancer

**DOI:** 10.64898/2026.05.04.720587

**Authors:** Elisa Gobbini, Pierre Duplouye, Marcelo Hurtado, Anne-Claire Doffin, Alexia Gazeu, Léo Hermet, Martial Scavino, Justine Berthet, Leo Laoubi, Sylvie Lantuejoul, Nicolas Gadot, Bertrand Dubois, Audrey Page, Eleonora Sosa Cuevas, Marie-Cécile Michallet, Laurent Greillier, Lionel Falchero, Jean Bernard Auliac, Marie Bernardi, Sophie Bayle, Marie Marcq, Julian Pinsolle, Stéphane Hominal, Olivier Bylicki, Sabine Vieillot, Fabrice Barlesi, Frédérique Penault-Llorca, Emmanuel Barillot, Stephane Depil, Margaux Hubert, Christophe Caux, Nicolas Girard, Vera Pancaldi, Jenny Valladeau-Guilemond

## Abstract

Dendritic cells (DCs) are central orchestrators of antitumor immunity. Several DC subsets—including conventional type 1 (cDC1), conventional type 2 (cDC2), plasmacytoid DCs (pDCs), and mature DC populations—play distinct roles in immune surveillance, tumor control, immunotherapy response and prognosis. Recent findings suggest that cDC1 are spatially closed to CD8⁺ T-cell and contribute to tertiary lymphoid structure formation in lung cancer. However, how other DC subsets interact with cDC1 to shape the tumor microenvironment (TME) remains largely unknown. Here, we analyzed the spatial distribution of major DC subsets, including cDC1, cDC2, mature DC and pDC, together with CD8⁺ T cells in a cohort of anti-PD1-treated NSCLC patients and we deciphered the corresponding immune microenvironment behavior by paired transcriptomic analysis. We found that, while other DC subsets populated the stroma, cDC2 were localized both in the stroma and in tumor nests. Moreover, unlike other DC subsets, cDC2 abundancy did not affect ICB response both at transcriptomic and *in situ* analysis. We described spatial organization of DCs in megaclusters characterized by distinct proportions of DC subsets. Patients enriched in megaclusters involving variable proportion of pDC, cDC1 and mature DC, exhibited pro-inflammatory transcriptomic programs while those enriched in cDC2-based megaclusters showed limited immune activation features. Globally, DC in lung cancer were structured around three distinct DC spatial patterns, namely cDC1-driven, cDC2-driven and DC-Scattered, each defined by unique compositions of DC megaclusters, immune features and pathways activation profiles. Among them, the cDC1-driven pattern was associated to prolonged anti-PD1 response in two independent cohorts.

## INTRODUCTION

Lung cancer remains the leading cause of cancer-related mortality worldwide, with non-small cell lung cancer (NSCLC) accounting for approximately 85% of all cases (1). Despite advances in targeted therapies and the advent of immune checkpoint inhibitors (ICIs), the 5-year overall survival rate is around 18%-20%, underscoring the need for improved understanding of tumor-immune interactions and more effective immunotherapeutic strategies (2,3).

Dendritic cells (DC), have a unique ability to control T cell function at tumor site through their ability to present tumor-associated antigens to T cells and to produce critical T cell differentiation cytokines and chemokines. The DC family contains many different subsets with various specific functions. Type 1 conventional dendritic cells (cDC1) play a central role in antitumor immunity (4). They uniquely express CLEC9A and XCR1, molecules involved in the uptake of necrotic material and cross-talk with T and natural killer (NK) cells, respectively (5–7). They are specialized for antigen cross-presentation, senses double-stranded RNA via TLR3 and produces type I and III interferons, IL-12, and chemokines such as CXCL9–10, which recruit and activate T cells (8,9). cDC1 can also support CD4⁺ T-cell and mostly drive a Th1 polarization (10). Conventional type 2 DCs (cDC2) represent a more heterogeneous population capable of driving diverse CD4 T helper responses, including the Th1, Th2, and Th17 pathways (11,12). cDC2 are key regulators of tumor immunity with context-dependent pro- and anti-tumor functions. They can migrate to tumor-draining lymph nodes to present tumor antigens to CD4⁺ T cells, and their intratumoral abundance has been linked to higher CD4⁺ T-cell infiltration in melanoma (11). Beyond direct antigen presentation, cDC2 can transfer antigens to lymph node–resident DCs and, similar to cDC1, cDC2 are capable of IFN-I–dependent cross-presentation to CD8⁺ T cells (13–15). However, some mouse studies suggest that CD11b+ cDC2 may also promote tumor progression by constraining antitumor CD4⁺ T-cell responses (16). Plasmacytoid DCs (pDCs) are defined by the expression of specific surface markers such as CD123 and BDCA2. In cancer, pDCs can play a dual role. Even though they may contribute to anti-tumor immunity via innate or adaptive responses (17), their production of type I interferons (IFN-I) can be suppressed by soluble factors like TGF-β and TNF-α in the tumor microenvironment (18,19), promoting the expansion of regulatory T cells (Tregs)(20). Finally, mature DCs, characterized by high expression of activation markers such as DC-LAMP/CD208 and CCR7, are consistently found across various cancer tissues (21). These cells can engage a crosstalk with T lymphocytes and NK cells through the production of IL-12, thereby supporting anti-tumor immune responses (22). Those mature DC detected in the T cell-rich areas of TLSs, adjacent to HEVs are often correlated to a good prognosis in many carcinomas including NSCLC (23)

Previous studies on transcriptomic data suggested a positive prognostic impact for pDC, cDC1 and cDC2 signature enrichment in lung cancer patients (9,24) but their performance as biomarkers is limited, suggesting that DC-related features other than density might be more revealing as markers of treatment response. Different DC subsets have been identified in lung cancer by scRNA sequencing and CITE-seq including pDC, cDC1, cDC2, DC3 and mature DC (22,25,26). The presence of these populations in lung cancer has also been confirmed by cytometry and immunohistochemistry, despite some controversy on the appropriateness of markers used for subsets identification. Indeed, while there are specific markers such as BDCA2 for pDC and CD208/DC-LAMP for mature DC (23,27,28), less-specific markers or negative selection strategies are usually needed to identify other populations (27,29). With these caveats, some studies have revealed that, in normal lung tissue, the majority of DCs are cDC2, with low numbers of DC3 and rare pDCs. Some DC expressing CD207/Langerin were described as Langerhans cell like (30,31) and may represent a cDC2 subset as they share many common marker with cDC2. In contrast, primary tumors exhibit similar levels of cDC2 and DC3 as well as increased pDC infiltration (26,32). Furthermore, cDC1s reduced in lung cancer relative to healthy tissue (25,26). Interestingly, cDC1 have recently been described as enhancing CD8⁺ T-cell activation and tertiary lymphoid structure formation in pre-clinical lung cancer models, supporting their central role in the tumor response, despite their low abundance (33). So far, no data are available confirming this compartmentalization of DC subsets at the tumor bed by using *in situ* approaches. Understanding the infiltration patterns and spatial distribution of DC subsets within NSCLC, as well as their impact on the surrounding tumor microenvironment, could provide valuable insights for the development of DC-based next generation treatments.

In this study, we examined the spatial organization of key DC subsets—cDC1, cDC2, pDC, and mature DC—alongside CD8⁺ T cells in a cohort of NSCLC patients treated with immune checkpoint blockade (ICB). All DC subsets were predominantly enriched in the stromal compartment, whereas cDC2 was present in both stroma and tumor nests. Notably, cDC2 density did not correlate with ICB response, in contrast to cDC1. We further delineated eight distinct DC spatial patterns, which we referred to as “megaclusters,” each defined by unique DC subset composition and spatial patterns and associated with specific immune microenvironments and pathway activations. Those megaclusters dominated by cDC2 and cDC1 suggest the presence of various immune niches, with cDC1-associated niches exhibiting the most robust predictive value across 2 independent cohorts.

## METHODS

### Cohorts of patients

The LYON-LUNG PREDICT_LATE cohort included 89 patients diagnosed with an advanced NSCLC and treated with anti-PD1 alone in first line setting. FFPE diagnostic samples and electronic medical records were collected thanks to the collaboration with two national academic societies namely, Groupe Francais de Pneumo-Cancerologie (GFPC) and UNICANCER. 267 samples were centralized at the Centre Léon Bérard Pathology Core and reviewed for quality validation in order to selected samples with adequate tissue amount This led to the selection of 60 patients for the analysis (**Figure 1A**). Each sample was used to prepare a 4 μm slide for multiplex immunofluorescence (mIF). Laser microdissection was then performed for RNA sequencing. Importantly, RNA-seq was conducted on serial slides corresponding to those used for mIF, targeting the same annotated region of interest. In order to validate our results, we used a second large cohort of advanced NSCLC patients (N = 122, CURIMMUNE) treated in first line setting as per standard of care with anti-PD1 alone or combined with chemotherapy. The CURIMMUNE cohort was provided by the CURIE INSTITUTE and we obtained access to both bulk RNAseq and electronic medical reports. The use of clinical samples and the collection of associated clinical data were conducted in compliance with French data protection regulations and have been declared to the CNIL in accordance with the MR-004 reference methodology (UNICANCER- SAFIR-02/CHECK’UP cohorts, declaration n°2221241 v0 of 18/02/2021; CURIMMUNE cohort, declaration n°2215984 v0 of 9/12/2019; GFPC-LUNGPREDICT-Late cohort, declaration n°2209524 v0 of 09/11/2018).

**Figure 1.**
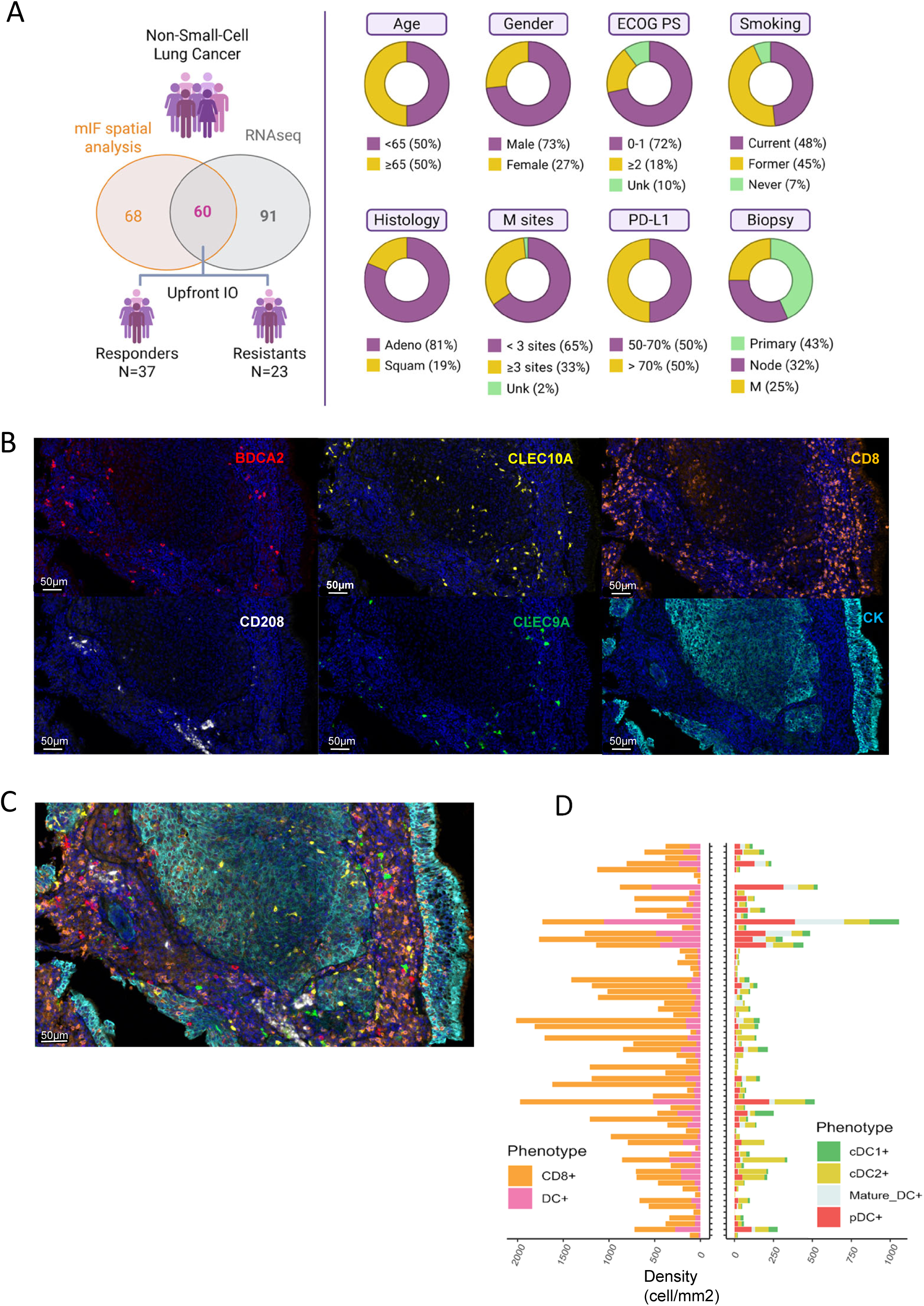
Detection of dendritic cell subsets in lung cancer tissues by in situ assays. (A) Lung Predict cohort overview. RNA sequencing and multiplex immunofluorescence (mIF) assays for dendritic cell detection were available for N = 60 patients. Clinical characteristics of the study population are presented. M: metastasis. **(B)** Representative single immunofluorescence images of lung cancer tissue showing cDC1 (CLEC9A), cDC2 (CLEC10A), pDC (BDCA2), mature DC (CD208), CD8⁺ T cells (CD8), and tumor cells (CK) staining. **(C)** Example of a multiplex immunofluorescence image of lung cancer tissue enabling simultaneous detection of cDC1 (green), cDC2 (yellow), pDC (red), mature DC (white), CD8⁺ T cells (orange), and tumor cells (turquoise). **(D)** Bar plots representing the densities of dendritic cell subsets and CD8⁺ T cells across the entire FFPE slide for each sample (N = 68).

### Multiplex immunofluorescent (mIF) staining

Markers for tumor-infiltrating DC identification were chosen based on the previous literature. In particular we multiplexed, BDCA2 (Goat Polyclonal, R&D Systems, AF1376) for pDC, CD208 (Rabbit 1010E1.01, Dendritics, DDX0191P) for mature DC, CLEC9A (Rabbit, abcam, ab223188) for cDC1, CLEC10A (Rabbit, OriGene, TA322508) for cDC2, CD8 (Mouse C8/144B, DAKO, M7103) for T lymphocytes (LT) and Cytokeratin (Mouse AE1/AE3, DAKO, M3515) for lung epithelial cancer cells. We set up a seven-colors mIF by sequential staining cycles using the tyramide-signal amplification (TSA)-based Opal seven-color Automation Research Detection Kit (Leica, DS9777) on the Bond RX automatic stainer (Leica Biosystems). Secondary antibodies used to amplify the signal included the Anti-Goat IgG HORSERADISH Peroxidase (HRP) Conjugated (Donkey Polyclonal, Life Technologies, A16005), the Anti-Rabbit IgG, HORSERADISH Peroxidase (HRP) Conjugated (Goat Polyclonal, Life Technologies, A1604) and the Anti-Rat IgG, HORSERADISH Peroxidase (HRP) Conjugated (Goat Polyclonal, Thermo Scientific, 31470)(34).

### Imaging analysis and cell quantification

Stained FFPE samples were scanned with the Vectra® Polaris™ Imaging System (Akoya, Biosciences, Marlborough, MA, USA) at 20X magnification. Slides were then visualized with Phenochart (1.0.12 version, AKOYA Biosciences) for multispectral imaging, annotated and quantified by inForm Tissue Analysis Software (Akoya, Biosciences, Marlborought, MA, USA). The whole tumor area including the invasive margin, defined as the region centered on the border separating the host tissue from the malignant nests with an extent of 1 mm, was analyzed for each sample. 26 to 30 training images of 1mm2 from 5 samples were used to build each algorithm. Tissue segmentation was optimized to define tumor, stroma and empty/artefact areas. Cell phenotyping was trained to identify phenotypes of interest based on staining intensities and shape of cells. Cell phenotyping was trained to recognize pDC (BDCA2+), cDC1 (CLEC9A+), cDC2 (CLEC10A+), mature DC (CD208+ CK-) and CD8 LT (CD8+). Tumor cells were defined as cytokeratin positive cells, while pneumocytes were CD208+ CK+ cells. All cells which did not fall into one of the previous labels were annotated as “Other” and may include other immune and stromal populations in the tumor microenvironment. Cells annotated as “Pneumocytes” or “Other” were excluded from the analysis to focus on the relationship between cancer cells, DC subtypes and CD8+LT.

The phenotyping output was manually validated through a slide-by-slide review using the Napari viewer, in which INFORM-identified phenotypes were directly overlaid onto the original QPTIFF scans. All resulting tables were manually curated using thresholding strategies tailored to each slide and each phenotype to reduce false positives and false negatives. For instance, for cDC2, only cells exhibiting strong CLEC10A expression were retained, CLEC10A low cells being excluded, even when INFORM had initially classified them as cDC2. Regions affected by artifacts that produced inconsistent annotations were excluded from further analysis. Phenotype densities were then quantified using the phenoptrReports 0.3.3 (Akoya) package in R, allowing for accurate density measurements within the specific regions previously delineated by INFORM.

### Spatial analysis

Using the SPIAT package (version 1.6.0) (35), we identified “spatial patterns” of spatially proximal cells using the identify neighborhoods function applied exclusively to the DC subset. This function applies the DBSCAN algorithm, for which we set a radius of 50 µm to define neighboring cells and require a minimum of 10 cells to form a cluster that we call “megacluster”. We also performed a Hierarchical clustering using the ward.D2 method. We scaled the megaclusters density (number of megacluster per tissue area) to compare samples relative to each feature (megacluster) and see which samples are enriched for specific features. For that, we calculated the mean and standard deviation of each megacluster across all samples (Z = (x - mean)/SD).

We also quantified both global and localized entropy between the “spatial patterns/megacluster” and LT_CD8⁺ cells. Global entropy was calculated using the calculated entropy function applied to the entire image. Localized entropy was assessed by dividing each slide into a spatial grid and computing entropy within each subdivision. Finally, for each slide, we determined the proportion of hotspots, defined as grid regions whose entropy exceeded the global entropy value.

### Cohort RNA sequencing

The LYON-LUNG PREDICT_LATE cohort RNA sequencing was performed at the Sequencing Core facility of CRCL on an Illumina NovaSeq machine with a paired-end protocol. Libraries were prepared with the RNAexome FFPE kit from Illumina. Raw sequencing reads were aligned on the human genome (GRCh38) with STAR (v2.7.8a), with the annotation of known genes from gencode v37. Gene expression was quantified using Salmon (1.4.0) and the annotation of protein coding genes from gencode v37.

The CURIEMMUNE cohort RNA sequencing was previously performed at the Sequencing Core facility of Institut Curie with the Illumina TruSeq RNA Access technology. The RNA-seq data were then processed with the Institut Curie RNA-seq pipeline v4.0.042(35). The raw bulk RNA-seq read counts were normalized with TPM (Transcripts Per Million) and log-transformed (i.e., log(X+1)). Patients were stratified in Responders (R) and Non-Responders (NR) according to a first-line PFS cut-off of 3 months to make them comparable with the LYON-LUNG PREDICT_LATE cohort.

### Pathway activity calculation

Log2(TPM + 1) counts were used to calculate pathway activities using the PROGENy database (36), a compendium of publicly available signaling perturbation experiments based on footprint genes to yield a common core of 14 signaling pathways. Pathways activities were calculated using the Multivariate Linear Model (MLM) from the package decoupleR (v2.9.7) (37). For the DC signatures, we use specific gene markers corresponding to each subset and gene set variation analysis (GSVA) (38) was performed using the Gaussian kernel method to estimate non-parametric enrichment scores for each gene set in each sample using the R package GSVA (v.2.4.1).

### Immune cell-type deconvolution

TPM (transcript per million) normalized counts were used to estimate cell-type proportions for lymphocytes (B, T and NK cells), myeloid cells (monocytes, macrophages and dendritic cells) as well as cancer, endothelial, eosinophils, plasma, myocytes, mast cells and cancer-associated fibroblasts (CAFs). Deconvolution was performed using the CIBERSORTx (39) method through the package omnideconv (v0.1.0) and using the LM22 signature (40).

### Estimation of immune response scores

Immune-scores were estimated on the Log2 (TPM + 1) normalized counts using the EasieR package (v1.4.0) (41) to generate immune profiles on a per sample basis. Immune-scores are calculated using gene sets that have been validated in different publications (see Table 1) as signatures to estimate certain hallmarks of the immune response.

**Table 1:** Immunoscores.

| Immunoscores | Genes | References |
| --- | --- | --- |
| Cytolytic activity (CYT) | "GZMA", "PRF1" | (63) |
| Roh immune score (Roh_IS) | "GZMA", "GZMB", "PRF1", "GNLY",<br>"HLA-A", "HLA-B", "HLA-C", "HLA-E",<br>"HLA-F", "HLA-G", "HLA-H", "HLA-DMA", "HLA-DMB", "HLA-DOA",<br>"HLA-DOB", "HLA-DPA1", "HLA-DPB1", "HLA-DQA1", "HLA-DQA2",<br>"HLA-DQB1", "HLA-DRA", "HLA-DRB1", "IFNG", "IFNGR1", "IFNGR2",<br>"IRF1", "STAT1", "PSMB9", "CCR5",<br>"CCL3", "CCL4",<br>"CCL5", "CXCL9", "CXCL10",<br>"CXCL11", "ICAM1", "ICAM2",<br>"ICAM3", "ICAM4", "ICAM5",<br>"VCAM1" | (64) |
| Chemokine signature (chemokines) | "CCL2", "CCL3", "CCL4", "CCL5",<br>"CCL8", "CCL18", "CCL19", "CCL21",<br>"CXCL9", "CXCL10", "CXCL11",<br>"CXCL13" | (65) |
| Davoli immune signature (Davoli_IS) | "CD247", "CD2", "CD3E", "GZMH",<br>"NKG7", "PRF1", "GZMK" | (66) |
| IFNy signature (IFNy) | "IFNG", "STAT1", "CXCL9", "CXCL10",<br>"IDO1", "HLA-DRA" | (67) |
| Expanded immune signature<br>(Ayers_expIS) | "GZMB", "GZMK", "CXCR6",<br>"CXCL10", "CXCL13", "CCL5",<br>"STAT1", "CD3D", "CD3E",<br>"CD2", "IL2RG", "NKG7", "HLA-E",<br>"CIITA", "HLA-DRA", "LAG3", "IDO1",<br>"TAGAP" | (67) |
| T-cell inflamed signature<br>(Tcell_inflamed) | "CCL5", "CD27", "CD274", "CD276",<br>"CD8A", "CMKLR1", "CXCL9",<br>"CXCR6", "HLA-DQA1",<br>"HLA-DRB1", "HLA-E", "IDO1",<br>"LAG3", "NKG7", "PDCD1LG2",<br>"PSMB10", "STAT1", "TIGIT" | (67) |
| Tertiary lymphoid structures<br>signature (TLS) | "CD79B", "CD1D", "CCR6", "LAT",<br>"SKAP1", "CETP", "EIF1AY", "RBP5",<br>"PTGDS" | (68) |

### Transcription factor activity inference and modules identification

Log2 (TPM + 1) counts were used to infer transcription factor (TF) activity. We use prior knowledge networks (PKN) to infer the activity of different TFs from the gene expression of its direct target genes quantified in the gene count matrix. We used CollecTRI a collection of transcriptional regulatory interactions, which provides regulons containing signed TFs - target gene interactions compiled from 12 different resources as database, accessed through the package decoupleR (v2.9.7)(37), and the Virtual Inference of Protein-activity by Enriched Regulon (VIPER) (v1.30.0) as the algorithm (42). Depending on the level of the counts and considering that one TF can have many targets and one target can be regulated by more than one TF, the algorithm can estimate the level of activity of the regulator based on correlation between target gene expression values.

For the analysis of the generated list of TFs, we adapted the weighted correlation network analysis (WGCNA) approach (v1.72-5) (8) to use it with TF activities and performed dimensionality reduction on these features by constructing what we defined as Weighted TFs co-activity networks (WTCNA) to detect highly correlated modules of TFs based on pairwise correlation of their inferred activity. Modules are defined as densely connected groups of nodes in the TF network, where connections represent correlation of activities, and they are arbitrarily named using colors. These TF modules were functionally characterized by calculating the Pearson correlation between these TF module scores and the pathways activity scores.

For context-specific differential TF activity, we inferred a regulatory network using ARACNe-AP (Algorithm for the Reconstruction of Accurate Cellular Networks with Adaptive Partitioning). ARACNe-AP reconstructs transcriptional regulatory networks by estimating mutual information between TFs and target genes and removing indirect interactions. The ARACNe-derived network was used to define context-specific TF regulons, which were subsequently employed for differential TF activity analysis (43,44).

Differential TF activity analysis is performed using msVIPER, a variant of VIPER that evaluates TF activity changes based on a differential expression-derived signature (42). To construct this differential signature, we apply a row-wise t-test comparing gene expression between conditions. The resulting p-values are converted into z-scores, which serve as input for msVIPER. Additionally, a null model is constructed using 1000 permutations of the dataset to estimate the background distribution of TF activity scores. Differentially activated TFs for each patient group are identified based on msVIPER enrichment scores, applying a significance threshold of pval<0.05. These TFs were then used to infer TF-signatures scores via GSVA for all the conditions.

### Survival analysis

We performed progression-free survival (PFS) analyses using custom R scripts built on the survival (v3.5.5) and survminer (v0.4.9) packages. Survival time and event status were first extracted to construct a Surv object. When predefined clinical groups were available, Kaplan–Meier curves were generated according to the specified grouping variable, and median PFS, 95% confidence intervals, and median follow-up (estimated using the reverse Kaplan–Meier method) were calculated. Group differences were assessed using the log-rank test. When no predefined groups were supplied, each feature was independently evaluated by assigning patients into high and low groups using a quantile-based threshold (median threshold = 0.5 by default) applied to the continuous feature values. Patients with feature values greater than or equal to the specified quantile were classified as “high,” while those below were classified as “low”. Kaplan–Meier curves and log-rank tests were then computed for each feature, and significant associations (p < 0.05 by default) were retained.

### Statistical analysis

Each patient’s response to treatment was assessed through progression free survival (PFS). PFS was defined as the duration from the initiation of first-line immunotherapy to the occurrence of the first progression event or last available status update, including the emergence of new lesions or the progression of pre-existing ones. Patients from the LYON LUNG PREDICT_LATE cohort were stratified in Responders (R) and Non-Responders (NR) according to a first-line PFS cut-off of 3 months for the analysis. Statistical analyses were performed using the Wilcoxon rank-sum test for comparisons between two independent groups. For comparisons involving more than two groups, one-way analysis of variance (ANOVA) was used, followed by Tukey’s honest significant difference (HSD) post hoc test for multiple comparisons. All graphs display individual samples with the median value indicated. Statistical significance is denoted as p < 0.05 (), p < 0.01 (), p < 0.001 (), and p < 0.0001 (****).

## RESULTS

### cDC2 infiltrates both stromal and cancer cell compartment while other DCs are mainly confined to the stroma

We analyzed formalin-fixed samples and clinical data from 60 NSCLC patients treated in first-line by pembrolizumab monotherapy as per standard of care (**Figure 1A**). In order to simultaneously analyze the spatial distribution of 4 DC subsets on whole tumors, we set up a novel multiplex immunofluorescence panel (**Figure 1B and 1C**). Notably, we used BDCA2 for pDC, CLEC9A for cDC1, and DC-LAMP/CD208 for mature DC identification. We coupled CD8 staining and Cytokeratin for detecting CD8 T lymphocyte (LT) and cancer cells respectively. Analysis of the transcriptomic profile of human DC from different lymphoid organs revealed the specific expression of CLEC10A marker by cDC2 together with the absence of cDC1 markers ((45) and **Figure S1A)**, despite its known expression on macrophages as well (46). After assessing CLEC10A and CD68 double staining, we observed that CD68⁺ cells exhibited lower CLEC10A intensity compared to CD68⁻ cells (**Figure S1B**). Therefore, we defined CLEC10A^high^ cells as bona fide cDC2 in our analysis. Slides were then scanned by Vectra® Polaris™ Imaging System, and analyzed with the inForm software. As shown in **Figure 1D**, the infiltration by CD8+ LT cells and DC subsets in NSCLC samples was remarkably heterogeneous. As expected, CD8+ LT cell infiltrated both the tumor nest and the surrounding stroma even though the CD8+ LT density was higher in the Stroma compartment (**Figure 2A**). Similarly, all DC subsets were enriched in the stroma surrounding the tumor compared to tumor nests (**Figure 2A**–**2B**). Interestingly, while pDC, cDC1 and mature DC were virtually absent in tumor nests, we unexpectedly found a relatively high median density of 9 cDC2/mm2 within the tumor islands (**Figure 2B**).

**Figure 2.**
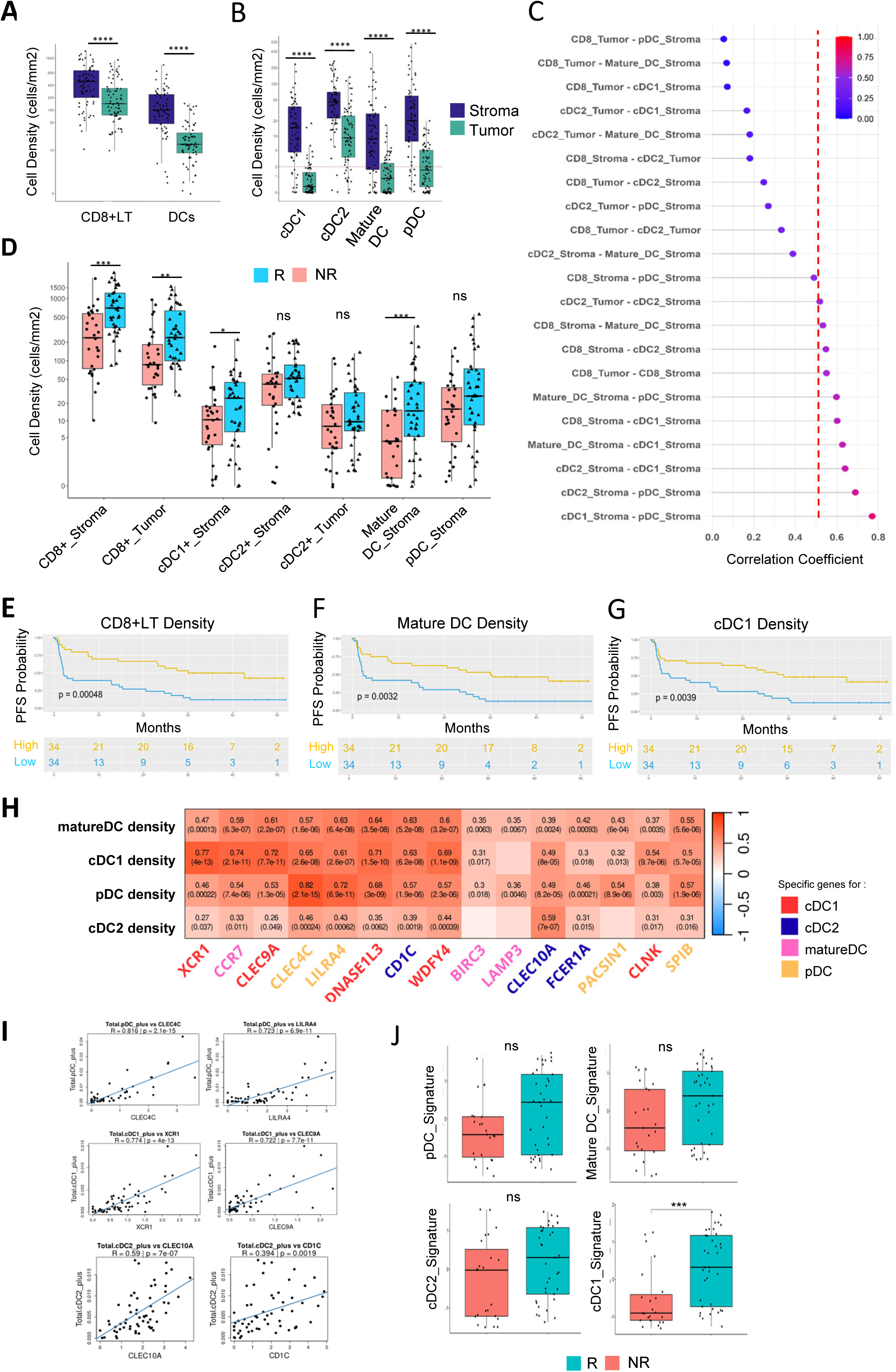
In contrast to cDC2, CD8⁺ T cells, mature DCs, and cDC1 are associated with ICB response. **(A)** Comparison of median nsities of CD8⁺ T cells and dendritic cells (DCs) in stromal versus tumor compartments across the entire study cohort. **(B)** Comparison of dian densities of DC subpopulations in stromal versus tumor compartments across the cohort. **(C)** Lollipop plot showing Spearman relation coefficients between immune cell populations detected in the stromal and tumor compartments. Coefficients are in ascending order m top to bottom. The red mark indicates the minimum R value (0.5) considered biologically meaningful. The color scale represents the ual R value. **(D)** Comparison of median densities of pDC, cDC1, cDC2, mature DC, and CD8⁺ T cells between responder **(n=38)** and non-ponder patients (n=30). **A-D,** Mann Whitney test; p ≤ 0.05 (*), p ≤ 0.01 (**), p ≤ 0.001 (***), p ≤ 0.0001 (****). **(E–G)** Kaplan–Meier curves for t-line progression-free survival (PFS) according to baseline densities of **(E)** CD8⁺ T cells, **(F)** mature DCs, and **(G)** cDC1. Patients were atified into “high” and “low” groups using the median density within the cohort as the threshold threshold (N=68 patients included for mIF). Heatmap showing Pearson correlation coefficients and their corresponding p-value in parenthesis between DC subset densities by mIF d expression of DC-specific genes by RNA-sequencing across the whole cohort (N=60 patients). The color scale represents the R value ly significant associations p-value < 0.05 are being shown). Genes are colored according to the DC subset they are specific for (red: cDC1; e: cDC2; pink: mature DC; yellow: pDC). **(I)** Scatter plot showing DC subset density (obtained in situ by mIF) with specific gene expression d linear regression**. (J)** Boxplots comparing the expression of DC-specific gene signatures between responders and non-responders were ted using one-way ANOVA. DC-specific gene signatures were calculated via GSVA using as gene sets the markers corresponding to each subset defined in H. Significant associations (p < 0.05) were followed by Tukey HSD post hoc tests. p ≤ 0.05 (*), p ≤ 0.01 (**), p ≤ 0.001), p ≤ 0.0001 (****).

In order to detected co-infiltration of different cell types, we analyzed the correlation between the density of each DC subsets and CD8 T cells. A modest correlation (r=0.5) was found between stroma- and tumor-infiltrating cDC2, suggesting that the cCD2 are not randomly distributed in the lung cancer tissue, but rather that different cDC2 subsets with various chemokine receptors and migration pathways may co-exist (**Figure 2C**). While stroma-infiltrating cDC1 and pDC showed the highest density correlation across samples (**Figure 2C)**, suggesting a particular cross-talk of these two DC subsets(47). Finally, we found that cDC1 showed a higher positive association with CD8⁺ T-cell infiltration compared to stroma-cDC2 or pDC (**Figure 2C**) suggesting a particular role of cDC1 to induce a CD8 T cell anti-tumoral response in human.

### In contrast to cDC1 and mature DC, pDC and cDC2, are not associated with immunotherapy response

We then investigated whether DC subsets enrichment was associated with response to immunotherapy. Patients were stratified into responders and non-responders according to their progression-free-survival (PFS) after first-line pembrolizumab calculated from the first injection of treatment. Responders were defined as having a PFS ≥ 3 months (62%) while non-responders had a PFS < 3 months (38%). Interestingly, patients responding to immunotherapy were enriched in stromal CD8⁺ T-cell, cDC1 or mature DC (**Figure 2D**). Even if not statistically significant, responders showed a numerically higher density of pDC as well. Conversely, cDC2 density was similar between responders and non-responders, regardless of their location in the stroma or in the tumor nest (**Figure 2D)**. Consistently, patients enriched in stromal CD8⁺ T-cell, cDC1 or mature DC had a longer PFS compared to the others (**Figure 2E-G).**

To confirm our observations at the transcriptomic level, we assessed the prognostic impact of DC subsets signature enrichment. First, we tested the correlation between the DC-specific gene expression in the transcriptomic data and the DC-subsets densities calculated in the immunofluorescence analysis (**Figure 2H**). Indeed, a poor correlation is often observed between transcriptomic and *in situ* analyses when characterizing immune cell populations, particularly when these subsets are rare and therefore more susceptible to sampling noise and detection limits (48). To improve the probability of achieving a robust correlation, both immunofluorescence and RNA sequencing were performed on the same FFPE block, and the *in situ* assessment was carried out precisely on the tissue area designated for microdissection on a subsequent slide. Interestingly, we found a good correlation between cDC1 specific genes expression such as CLEC9A and XCR1 and the cDC1 density (**Figure 2H-I**; r= 0.77 and 0.72 respectively). Same result is obtained with pDC density with a correlation coefficient of 0.82 and 0.72 with CLEC4C and LILRA4 respectively, and thus demonstrating the feasibility of using the expression of these specific genes to quantify the infiltration of these two populations in tumor bulk RNAseq data. In this context, we found a good correlation between specific DC subset signature expression as defined in Table 2 and their measured tissue densities in the study cohort (**Figure S1C-D**), notably for pDC and cDC1 (r=0.66 and 0.69 spearman correlation coefficient respectively). Mature DCs showed the lowest correlation coefficient (r = 0.48, p =9.6 10^-5^) between signature expression and *in situ* cell density, consistent with the potentially lower stability of transcripts such as *BIRC3* and *LAMP3*, whose expression may reflect a transient activation program rather than a stable ontogenetic lineage.

**Table 2:**
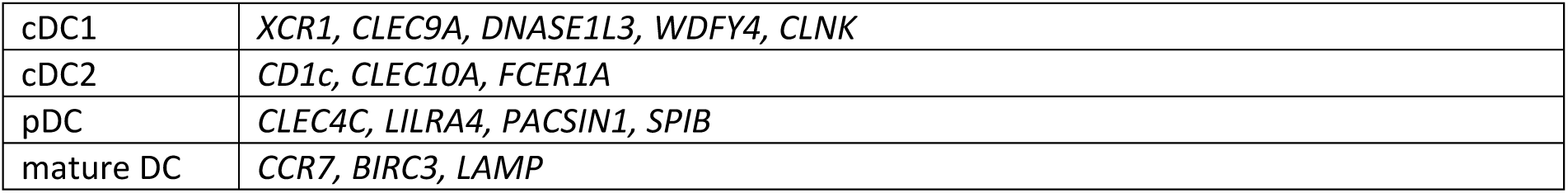
DC signatures.

Using these DC specific signatures, we confirmed the positive prognostic impact of the cDC1 infiltration in lung cancer by comparing the cDC1 signature between responders and non-responders. Although not statistically significant, responders were also enriched in mature DC and pDC. Conversely, cDC2 signature enrichment was similar across responders and non-responders confirming our immunofluorescence results (**Figure 2J**).

Overall, these results confirmed the positive prognostic impact of stromal cDC1 in lung cancer patients treated with immunotherapy. Results for mature DC and pDC were less consistent, but a tendency for a positive prognostic impact was highlighted. Conversely, we demonstrated that cDC2 had no impact on immunotherapy response, confirming a distinct role for this subset in the anti-tumor immune response.

### DC subsets are structured in megaclusters with different consequences on immune responses

To investigate the relationships among DC subsets and how they interact to elicit immune responses, we performed a spatial analysis using annotated DCs. The spatial coordinates of each DC, obtained through *in situ* analysis, were used to characterize their organization in megaclusters within the cancer-associated stroma (**Figure 3A)**. Neighbourhood analysis enabled the identification of 4529 megaclusters across the entire cohort, representing spatially localized groups of DC, each characterized by a distinct composition of DC subsets (**Figure 3B**). Using a hierarchical clustering approach, we further defined eight classes of megaclusters shared across patients, each exhibiting a unique profile of DC subset proportion (**Figure 3B**–**3C**). Within these 8 classes, we identified 4 megaclusters profiles mainly composed by a given DC subset, namely Enriched_cDC1 (12.89% of the 4529 clusters), Enriched_cDC2 (12.39%), Enriched_mature DC (17.60%) and Enriched_pDC (34.93%) (**Figure 3C**–**3D**). Moreover, we found some of these to fall into mixed profiles namely Mixed_cDC1_pDC (6.40%), Mixed_pDC_mature_DC (5.45%), Mixed_cDC2_pDC (7.64%) and Mixed_cDC2_mature_DC (2.69%). Interestingly, we did not find megaclusters with a mixed population of cDC1 and cDC2 suggesting the absence of interaction between these two conventional DC subsets and possibly different anti-tumor functions. No mixed cDC1/mature DC megaclusters were found, suggesting that either, once activated, mature cDC1 lose their subset-specific marker and acquire maturation markers either they migrate in a different area to perform their function. If we compare the density of each megacluster in responders versus non-responder patients, none of them was statistically different and thus not associated with response to immunotherapy (**Figure S1E**). Nevertheless, some mixed megaclusters—namely Mixed_cDC1_pDC, Mixed_cDC2_mature_DC and Mixed_pDC_mature DC—showed a trend toward higher density in responders compared to non-responders.

**Figure 3.**
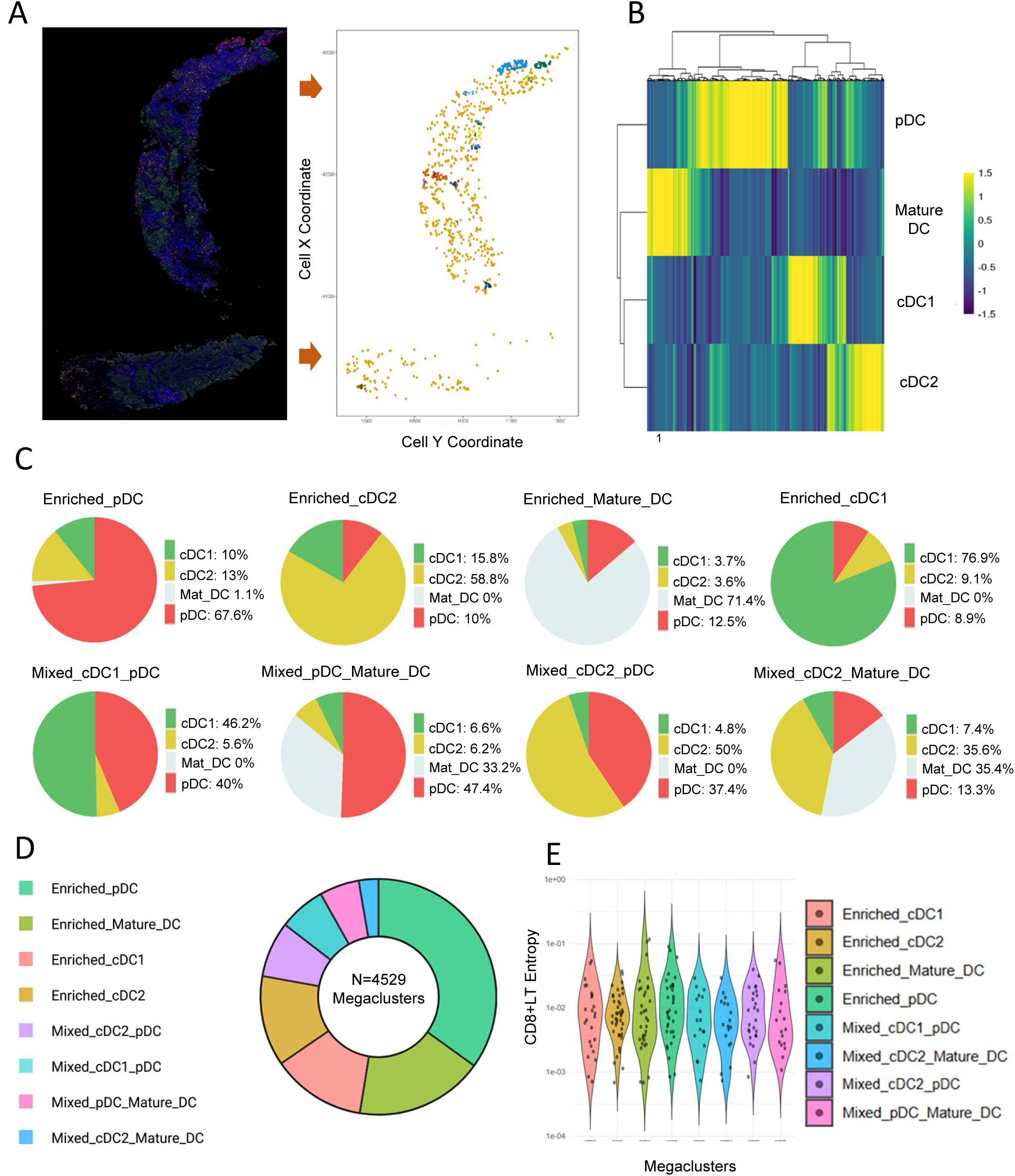
DC subsets are organized into 8 spatial niches with similar CD8+LT density. **(A)** Representative example of a multiplex immunofluorescence staining and the corresponding 2D spatial map of annotated dendritic cells (DCs) after image analysis. Colored dots represent individual DCs belonging to the same megacluster; DCs sharing the same color are part of the same megacluster. In this example, 13 megaclusters were identified. **(B)** Hierarchical clustering (ward.D2) of DC megaclusters based on the relative proportions of DC subsets within each megacluster, revealing 8 distinct spatial niches (megacluster types) recurrently detected across patients. The color scale represents the relative enrichment (z-score or normalized proportion) of each DC subset within each megacluster.**(C)** Schematic representation of the composition of each megacluster type, showing the proportion of included DC subsets.**(D)** Overall distribution of the 8 megacluster types within the total pool of identified megaclusters. **(E)** CD8+ lymphocyte (CD8+LT) density (entropy) across the 8 megacluster types, highlighting comparable CD8+ LT infiltration among them (n=68 patients).

In order to investigate the functional properties of megacluster classes in the stroma, we analyzed the CD8⁺ T-cell entropy as a proxy of CD8⁺ T-cell density in each megacluster (**Figure 3E).** CD8⁺ T-cell entropy was comparable across megacluster classes, suggesting that the functional specificities of megacluster classes are not primarily driven by direct physical interactions of DCs with LT.

In order to explore some functional specificities linked to DC density, we integrated the transcriptomic features with the megaclusters densities defined by the *in-situ* analysis. By using correlation networks of transcription factor (TF) activity, we first identified ten TF modules describing pathway activation status in each tumor sample (denoted by colors, see Table 3; **Figure 4A**). We focused our analysis on the magenta, purple, black, and yellow TF modules, as these displayed strong correlation coefficient with some TF enabling biological interpretation. The magenta module is characterized by high NF-κB and JAK/STAT signaling, consistent with a strong inflammatory and activated state within the tumor microenvironment. The black module shows a very similar profile, also enriched for NF-κB, JAK/STAT, and PI3K pathways, suggesting that these two modules capture closely related inflammatory and survival programs, likely reflecting overlapping tumor-intrinsic and microenvironment-driven signals. In contrast, the purple module is more specifically associated with JAK/STAT signaling without concomitant activation of broader stress or inflammatory pathways, and TRAIL-mediated cancer cells apoptosis, suggestive of anti-tumor immune processes distinct from broader inflammatory stress responses. Finally, the yellow module is distinguished by its association with PI3K, TGFβ, and hormonal signaling, suggesting a more differentiated state and an immune cold microenvironment.

**Figure 4.**
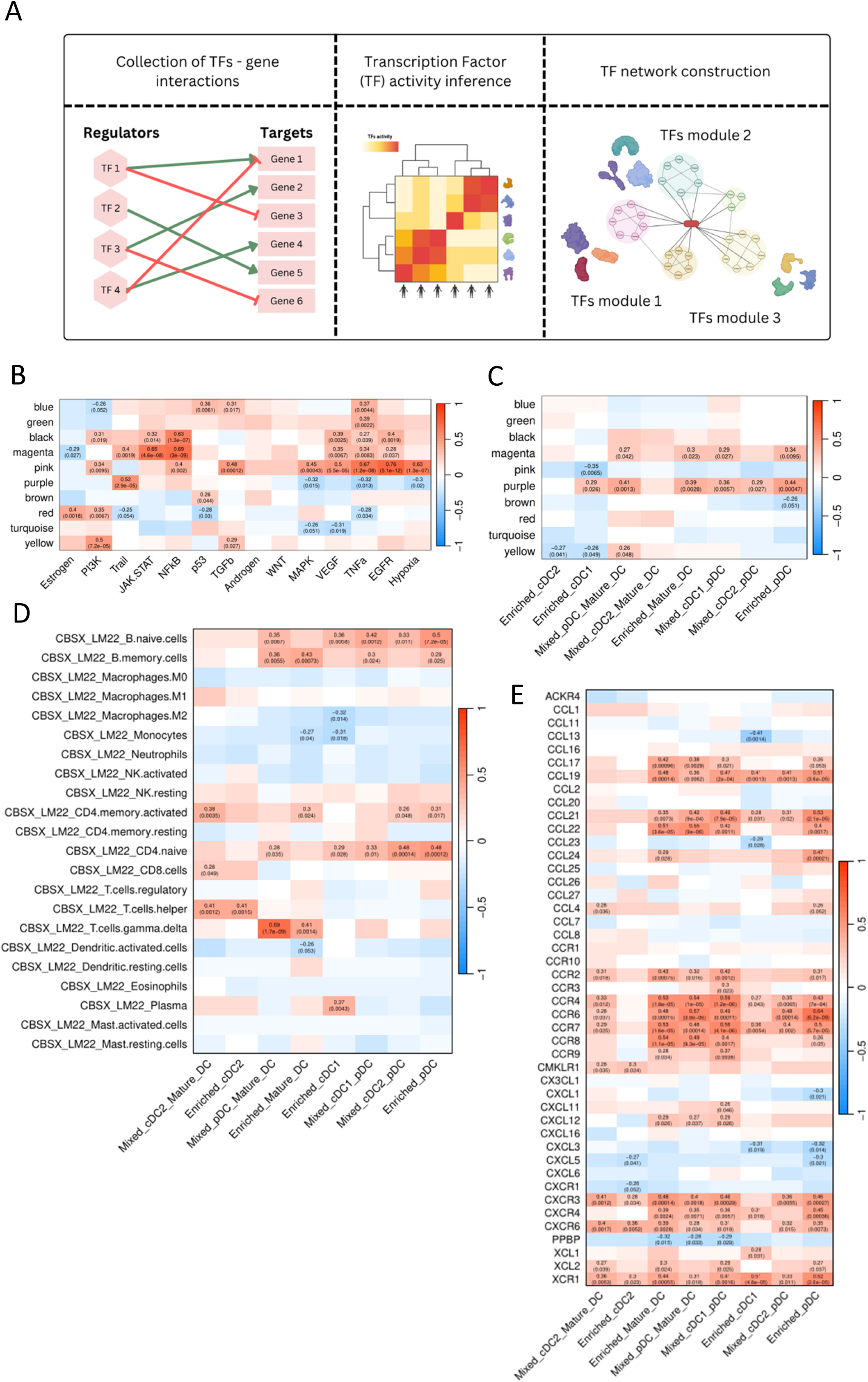
DC subsets are structured in megaclusters with different consequences on immune response. **(A)** Schematic representation of the workflow used for the identification of transcription factor (TF) modules from RNA-sequencing data.(B) Heatmap showing correlation coefficients between pathway activities and TF module activity across the study cohort. **(C)** Heatmap displaying the correlations between the TF modules and the density of megaclusters defined by multiplex immunofluorescence (mIF) analysis in the study cohort**. (D)** Heatmap of correlation coefficients between immune cell deconvolution scores (derived from CIBERSORTx and LM22 signature) and megacluster densities**. (E)** Heatmap showing correlations between cytokine gene expression and megacluster densities. In all heatmaps, the color scale represents the Pearson correlation coefficient (R value), and p-values are reported within each cell (only significant associations p-value < 0.05 are being shown).

**Table 3:**
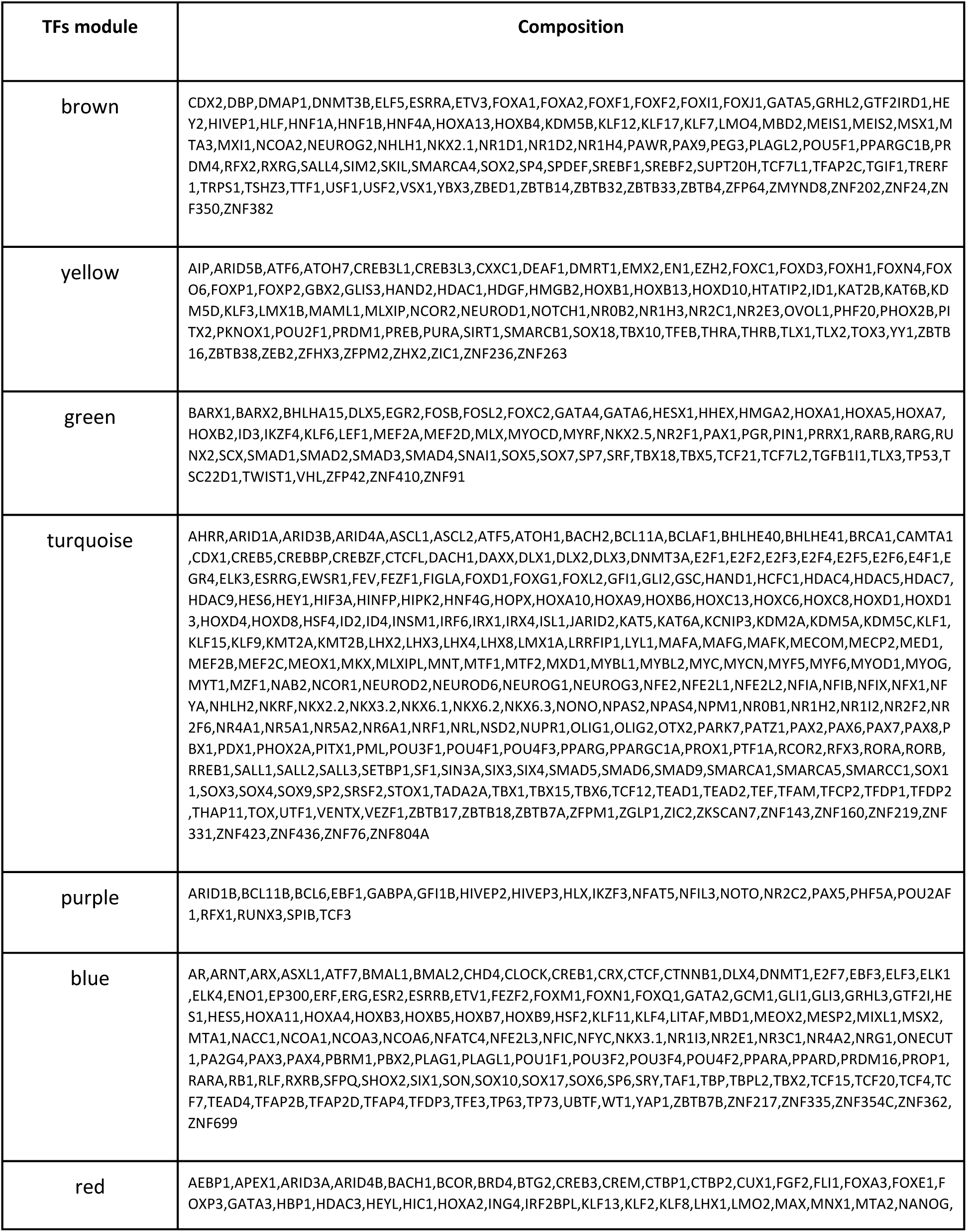
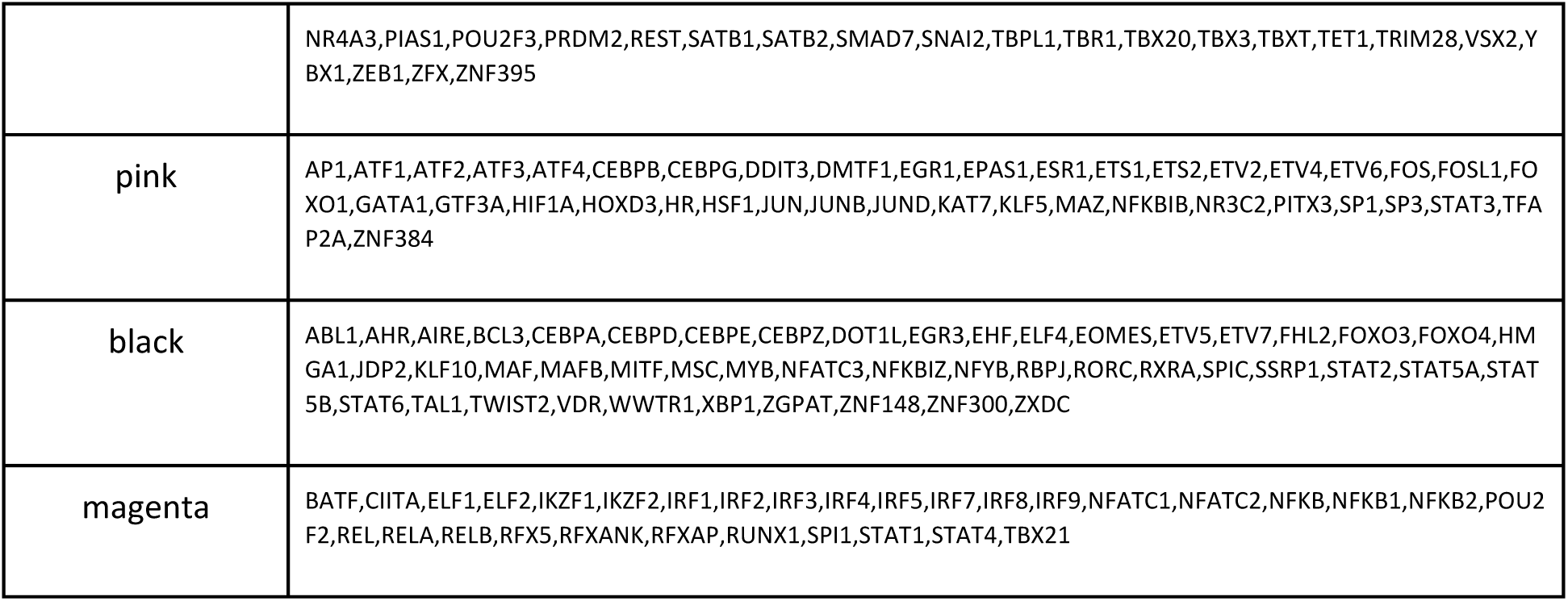
TF modules composition.

We then studied the correlation between TF module enrichment and megaclusters densities in the study population. Interestingly, the Purple and Magenta TF modules were broadly positively correlated with almost all megaclusters containing pDC, suggesting a central role of this DC subset in the immune response (**Figure 4C**).

In order to better decipher how megaclusters may be associated with tumor microenvironment composition, we then performed cell-type deconvolution analysis on the bulk transcriptomic data and investigated the correlation of different cell types and expression of specific genes with megaclusters densities (**Figure 4D-E**). Enriched_pDC megaclusters and most of mixed megaclusters containing pDCs were positively correlated with B-cell (naïve and memory) and CD4⁺ memory T-cell signature (**Figure 4D and suppl. Figure S2**). Consistently, we found a positive correlation of enriched_pDCs megaclusters and other pDC-centered megaclusters with gene expression of CCL19, CCL21, CCR7, and CCR6, suggesting their involvement in immune cell recruitment within the tumor microenvironment (**Figure 4E**). Notably, megaclusters that included cDC2 rarely showed enrichment in either the Purple or Magenta TF modules (**Figure 4C**). Consistently, deconvolution analysis revealed that the Enriched_cDC2 megacluster was poorly correlated with most immune cell types and gene expression of chemokines/chemokines receptors (**Figure 4D**–**4E)**. Of note, the only chemokine receptor associated with Enriched_cDC1 megacluster density is XCR1, in line with the preferential expression of the marker by cDC1 (**Figure 4E**). Finally, a high correlation was observed between the density of mixed_pDC_mature_DC megacluster and the γδ T cell signature but was due to one particular outlier (**Suppl. Figure S2**). Altogether, these results demonstrate that DCs are spatially organized into megaclusters characterized by distinct proportions of DC subsets. Megaclusters containing pDCs-enriched patients were associated with tumors with the most pronounced pro-inflammatory signatures. Conversely, cDC2-enriched megaclusters-enriched patients exhibited limited immune activation features in the tumor microenvironment, suggesting a limited contribution of this subset to the anti-tumor response.

### cDC1-driven spatial pattern is correlated to the response to immunotherapy

Given that the enrichment of individual DC megaclusters was not sufficient to predict immunotherapy response (**Supplementary Figure S1E**), we hypothesized that the co-occurrence of different megaclusters within the tumor microenvironment plays a more critical role in shaping effective antitumor immunity. We thus performed a hierarchical clustering on patients according to the density of the 8 megaclusters previously identified by the *in situ* approach (**Figure 5A and Suppl. S3A**). We identified three distinct patient groups, referred to as DC-scattered, cDC2-driven, and cDC1-driven, based on the prevalent DC subsets in the dominating megaclusters for each patient group. The DC-scattered group was characterized by a low proportion of all megaclusters, reflecting a sparse or mixed DC component **(Suppl. Figure S3B**). The cDC2-driven group primarily included patients enriched in cDC2-based megaclusters, such as Enriched_cDC2, Mixed_cDC2_mature_DC, and Mixed_cDC2_pDC. In contrast, the cDC1-driven group comprised patients enriched in cDC1-associated megaclusters, including Enriched_cDC1, Mixed_cDC1_pDC, and Mixed_pDC_mature DC. We then studied clinical characteristics of each of these 3 patient groups. The cDC1-driven group exhibited the most favorable PFS under first-line pembrolizumab in the LYON-LUNG PREDICT_LATE cohort, followed by the cDC2-driven group, whereas the DC-scattered group displayed the poorest outcome (**Figure 5B**). We next analyzed the corresponding transcriptomic data from patients classified as DC-scattered, cDC2-driven and cDC1-driven in order to identify differentially active TFs associated with each DC spatial pattern (see Methods). We thus generated a TF activity signature capable of classifying the three patient groups starting from bulk transcriptomic data (Table 4). As expected, the resulting TF signatures successfully recapitulated the prognostic impact of DC spatial organization patterns in the LYON-LUNG PREDICT_LATE cohort (**Figure 5C).**

**Figure 5.**
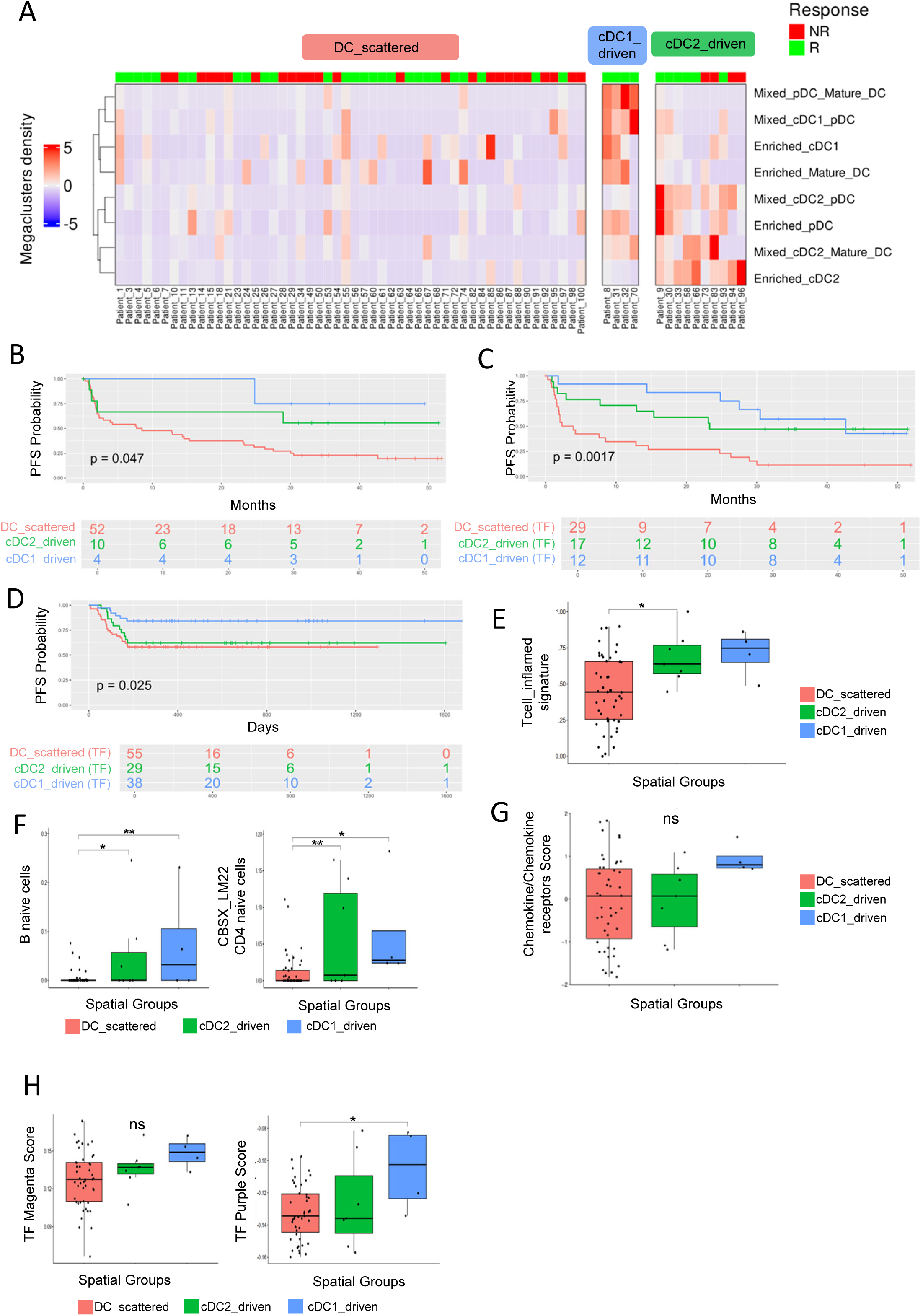
cDC1-driven pattern sustains a TLS-like inflammatory microenvironment. **(A)** Hierarchical clustering (ward.D2) of patients (N = 66) based on the density of DC-based megaclusters. The color scale represents megacluster density. Three patient groups were identified: DC-scattered, cDC2-driven, and cDC1-driven (66 patients (68 mIF patients - 2 patients who fall apart from the 3 main clusters - see Figure S2E). **(B)** First-line progression-free survival (PFS) Kaplan–Meier (KM) plot according to DC-driven spatial organization, as defined by the immunofluorescence assay in the study cohort. **(C)** First-line PFS KM plot according to the transcription factor (TF) signature corresponding to DC-driven spatial organization in RNA sequencing data from the study cohort (n=58 patients (60 patients from overlap RNAseq and mIF - 2 patients who fall apart from the 3 main clusters - see Figure S2E). **(D)** First-line PFS KM plot according to the TF signature corresponding to DC-driven spatial organization in the validation cohort (CURIEIMMUNE, N = 122). **(E)** Boxplot showing T cell–inflamed signature expression across spatial groups. **(F)** Boxplots showing expression of B naïve cell (left) and CD4 naïve T-cell (right) deconvolved signatures (CIBERSORTx_LM22) across spatial groups. **(G)** Boxplot of chemokine and chemokine receptor signature expression across spatial groups. **(H)** Boxplots showing expression of TF magenta (left) and TF purple (right) modules across spatial groups (N=58). For all boxplots, statistical significance was assessed using one-way ANOVA, followed by Tukey’s honest significant difference (HSD) post hoc test for multiple comparisons: p ≤ 0.05 (*), p ≤ 0.01 (**), p ≤ 0.001 (***), p ≤ 0.0001 (****).

**Table 4:**
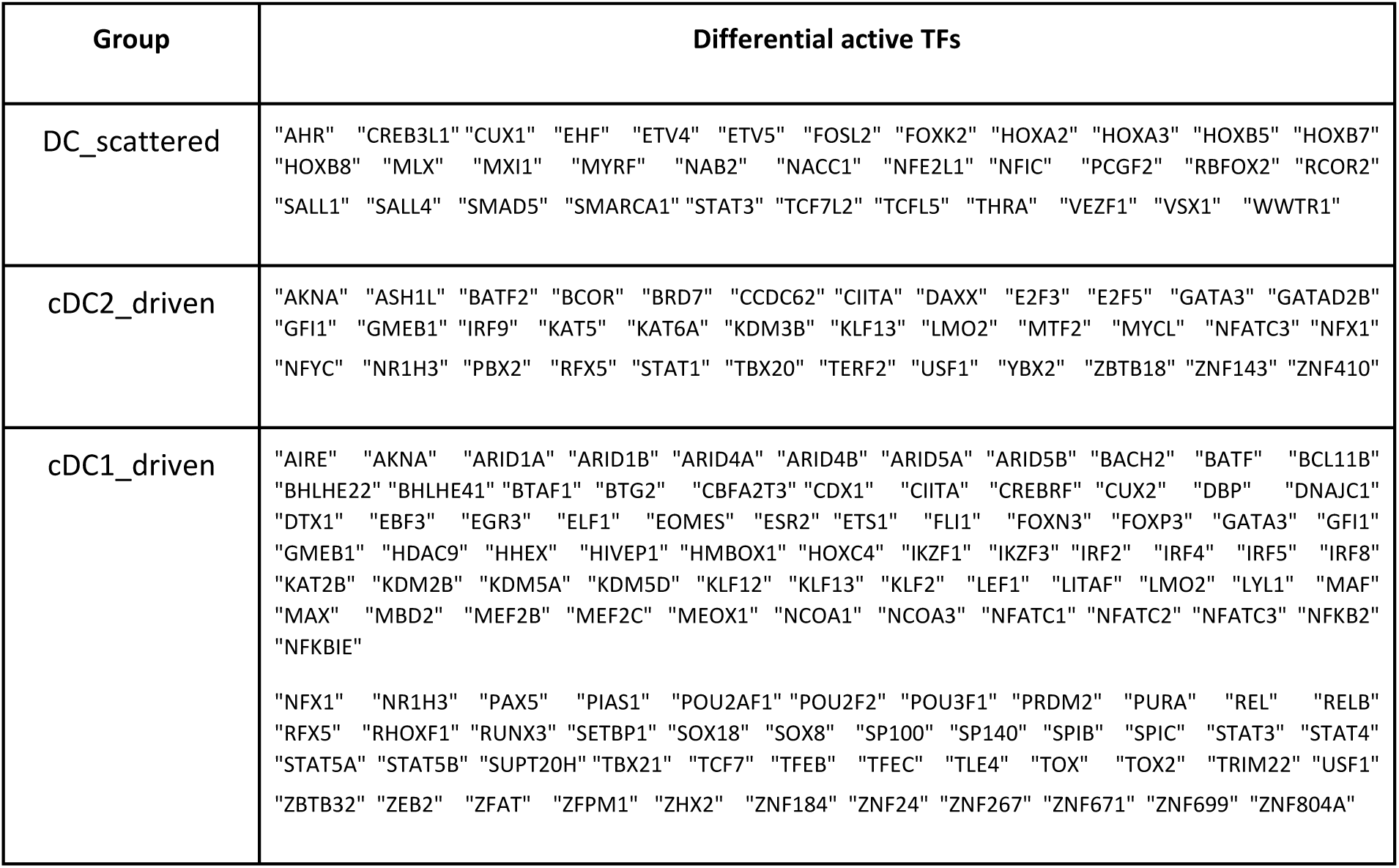
Differential active TFs.

Having identified a transcription-derived signature for the three DC organization patterns we applied this method to validate our results in an independent cohort with already available transcriptomic data (CURIEIMMUNE cohort; see Methods). The Principal Component Analysis (PCA) performed on transcriptomic data from the study cohort and the validation cohort revealed a pronounced batch effect when all transcriptomes were taken into account (**Figure S3C left panel**). However, this effect was absent when restricting the analysis to TF activities (**Figure S3C right panel**). The TF-based cDC1-driven group was confirmed to achieve the better PFS compared to the other ones in the CURIIMMUNE cohort as well supporting the robustness of our results (**Figure 5D**).

To investigate how the global spatial organization of DCs influences the tumor-associated stroma, we analyzed the transcriptomic features of the three DC spatial patterns defined according to the *in situ* analysis in the LYON-LUNG PREDICT_LATE cohort, with a particular focus on the immune compartment. cDC1-driven samples exhibited the strongest enrichment in T-cell inflamed signature (**Figure 5E**). The deconvolution analysis further revealed that cDC1-driven patients showed the highest B naïve and naïve CD4⁺ T cells enrichment (**Figure 5F**). Other immune cell populations were similarly enriched across groups (**suppl. Figure S3D)**. Although not statistically significant, cDC1-driven samples exhibited the highest expression of chemokine and chemokine receptor gene signature (**Figure 5G**). Among them, we found some key chemokine/chemokines receptor axes having a central role in TLS-like structures formation, such as CCL21, CCR4, CCL22, CCR6 and CCR7 (**Figure S3E**). Moreover, XCR1, a marker of cDC1 that mediates cross-talk with CD8⁺ T cells and facilitates antigen presentation in lymphoid-like niches, was also enriched in the same group compared to others. Finally, the cDC1-driven patient subgroup showed the highest enrichment for the pro-inflammatory TF modules (TFs purple and magenta; **Figure 5H**), suggesting a chronic immune engagement compared to the cDC2-driven group.

Overall, these results suggest that the DC spatial patterns are critical determinants of immunotherapy response, with cross-talk among pDCs, mature DCs, and cDC1s driving effective anti-tumor immunity. Again, cDC2s appear to play a limited role in reshaping the TME and sustaining the immune response.

## Discussion

cDC1 participate in cross-talk with key players in the anti-cancer response such as CD8+ T cells, NK cells and CD4+ T cells, being able to elicit effective cancer recognition by the immune compartment (17). Consequently, cancer activates several mechanisms to impair cDC1 function and development in order to facilitate progression. For instance, cancer cells can produce prostaglandin E2 and IL-6 which inhibit late-stage cDC1 function and hinder the differentiation of cDC1 progenitors, respectively (49,50). Moreover, tumor-secreted gelsolin inhibits CLEC9A-dependent cross-presentation of dead cell-associated antigens by cDC1 (51). Finally, in lung cancer mouse models, cDC1s can be dysfunctional because of downregulation of antigen uptake receptors such as TIM4, diminished cross-presentation and decreased expression of adhesion molecules such as ALCAM to interact with cytotoxic T cells.(52,53). Given the central role of cDC1 in cancer recognition, preclinical models have suggested that the absence of cDC1, negatively impacts the anti-PD-L1 immunotherapy response (54,55). In a conditional knock-out model for PD-L1 suppression on cDC1, inhibition of PDL-1 on cDC1 was sufficient and necessary to inhibit tumor growth increasing the infiltration of CD8+ T cells with an exhausted phenotype (PD1+, LAG-3+, TIM-3+) into the TME (56). Consistently, high tumor-infiltrating cDC1 density and cDC1-related signature are associated with better prognosis and favorable response to immune checkpoint inhibitors in several human cancers (7,9,57,58). We previously showed that the cDC1-signature is associated with improved overall survival in lung cancer as well, however, a validation by *in situ* quantification of tumor-infiltrating cDC1 has so far been lacking. Here we showed for the first time that, although tumor-associated cDC1 are relatively rare compared to other DC subsets such as cDC2 or pDC, they are strongly associated with immunotherapy response in lung cancer patients. Moreover, according to our findings, the prognostic value of cDC1 infiltration, is much stronger than pDC or mature DC densities suggesting a more direct and non-redundant role of cDC1 in anti-cancer response. We previously identified the privileged interaction between cDC1 and CD8+ T cells in melanoma patients by addressing the spatial distribution of immune cells within the tumor bed. Here we found that cDC1 was the only subset showing a statistically significant positive association with CD8⁺ T-cell infiltration in lung cancer. The lack of correlation between CD8⁺ T-cell density and the infiltration of other DC subsets suggests that other DC subpopulations may orchestrate a broader immune response beyond the activation of CD8⁺ T lymphocytes which is mainly linked to cDC1. Moreover, our spatial analysis provided previously unreported insights into how cDC1 interacts with other DC subsets for mediating their functions. Stroma-infiltrating cDC1 and pDC showed the highest density correlation across samples and could cluster together in a spatially organized niches as we previously described in melanoma (59). Proficient cross-talk between pDC and cDC1 has been suggested by preclinical studies in mouse models. For instance, the immunization of E.G7 inoculated mouse model with antigen-pulsed pDC and/or cDC showed that pDC can enhance the capacity of cDC to activate antigen-specific CD8+ T cells and limit tumor growth (60). Recently, we demonstrated that, in human cancers, cDC1 promotes pDC survival and restores IFN-α production counteracting the immunosuppression mediated by TGF-β or PGE2 in the TME through the release of IFN-III (47). IFN-I producing pDC play a central role in Tertiary Lymphoid Structures (TLS) formation and function in lung cancer (61) and have also been identified within TLS in colon cancer patients. Multiplex immunofluorescence revealed the presence of pDC in the T zone, in close proximity to CD4+ T cells. Consistently, our transcriptomic analysis revealed that the presence of pDC niches is associated with a TME composition and cytokine expression profile indicative of a lymphoid-like organization. Moreover, we detected a pDC-mature DC niche supporting spatial proximity between pDC and mature DC which are usually localized within TLS in the lung cancer context (23,62). On the other hand, cDC1 have been recently suggested to play a central role into TLS formation in lung cancer preclinical models(33). Interestingly, no mixed cDC1/mature DC niches were found in our analysis, suggesting that once activated, mature cDC1 might lose their subset-specific marker, acquire maturation markers and migrate into the TLS as mature DC. Several studies have shown the positive prognostic impact of TLS in lung cancer and patients with higher TLS density have been showed to achieve a better response to immunotherapy (28). Consistently, our results revealed that, patients presenting a transcriptomic profile indicative of a cDC1-driven spatial pattern (characterized by close contact between cDC1, pDC and mature DC), achieved longer progression-free survival when treated with first-line anti-PD1 pembrolizumab across two independent cohorts.

Compared with cDC1, the functional properties of cDC2 in cancer are poorly understood. Previous studies conducted by scRNAseq and CyTOF demonstrated that cDC2 represent the most abundant DC subset in normal lung tissue (4). Here, we provided the first evidences of *in situ* localization of cDC2 in lung cancer, showing that while other DC subsets are virtually absent within tumor nests, cDC2 are well represented across both tumor and stroma compartments. These results support previous data regarding the physiological localization of cDC2 in lung tissue prior to malignant transformation. However, a modest correlation was found between stroma- and tumor-infiltrating cDC2, suggesting that the cDC2 are not randomly distributed in lung cancer tissue. Rather, distinct cDC2 subsets with different chemokine receptors and migration pathways may co-exist, consistent with the high heterogeneity of this population. For example, CLEC10A positive cells in tumor islets may have a more LC like phenotype as those cells were clearly also observed in tumor nest and will need further characterization. Previous studies suggested also a poor prognosis for cDC2-enriched patients with lung cancer and melanoma, however, those studies defined cDC2 based on BDCA1 expression, which is shared by other immune cells, limiting interpretation. Among others, we previously showed that enrichment of a cDC2 signature may confer improved overall survival in several malignancies, including lung cancer. Nevertheless, the limited number of genes included in these transcriptomic signatures cannot fully capture the complexity of such a heterogeneous population. Moreover, little is known about the impact of cDC2 on immunotherapy-treated patients. Here, we provided for the first time the evidence that, unlike other DC subsets, cDC2 infiltration has no impact on immunotherapy response. Consistently, we found no correlation between cDC2 and CD8⁺ T-cell infiltration. Similarly, cDC2-enriched niches were poorly associated with immune activation features, immune cell enrichment and cytokine gene expression in the TME, suggesting a limited contribution of this subset to anti-tumor immunity. While we identified DC niches supporting a potential cross-talk between cDC2 and pDC or mature DC, we did not detect megaclusters containing both cDC1 and cDC2 populations suggesting the absence of interplay between these two conventional DC subsets and distinct anti-tumor functions for each subset. Even so, while potential interactions between cDC2 and other DC subsets warrant further validation, the dichotomy between cDC1- and cDC2-driven spatial DC patterns was associated with distinct impacts on immunotherapy response. Patients presenting a cDC2-driven spatial pattern, experienced a shorter progression free survival under pembrolizumab compared with those displaying a cDC1-driven pattern. It remains to be elucidated whether these two DC spatial patterns represent different stages of the immune response activation and whether cancer treatments can modulate transition between them.

Among the limitations of this study, we acknowledge the lack of *in situ* validation of the spatial relationship between TLS and DC subsets suggested by transcriptomic analysis. Indeed, correlations between bulk RNA sequencing transcriptomic features and of DC niches enrichment defined by *in situ* approaches may introduced interpretation bias, which should be addressed using spatial transcriptomics in future studies. Such approaches would also help dissect cDC2 heterogeneity, providing insights into spatial relationships between cDC2 subsets and tumor compartments and clarifying their functional properties. Moreover, due to the lack of specific biomarkers, we could not explore the spatial distribution of DC3, whose role in cancer biology remains to be clarified.

Despite these limitations, here we provided the first evidence of spatial organization of major DC subsets in a large cohort of lung cancer patients, including cDC2, which had not been previously described. Paired transcriptomic analyses, allowed identification of TME immunological features associated with distinct DC spatial patterns, highlighting cDC1-driven profiles as the most favorable for immunotherapy response. In contrast, our results show that cDC2 exert a more limited effect on TME remodeling and immunotherapy response. Altogether, these findings confirm cDC1 as the most promising DC subset for targeted therapies aimed at remodeling the TME and triggering effective anti-tumor immune responses.

## Contribution

EG collected samples as Principal Investigator of the LUNG PREDICT_LATE cohort, she designed experiments, she performed along with A-CD the multiplex staining and the Inform analysis, she analyzed results and she wrote the paper. PD performed the bioinformatics and statistical analysis on the multiplex in situ data and he revised the manuscript. MHurtado performed the bioinformatics and statistical analysis on the RNAseq data and the integration with imaging features. L.H. and M. S. performed controls on image analysis. J.B. provided advices for in situ staining. AG, SL and NG performed the pathological annotations. LG, LF, J-BA, MB, SB, MM, JP, SH, OB, SV, FB and FP-L provided the samples collected in the LUNG PREDICT-Late cohort. E. SC, A. P., S.D., M-CM., C.C. and B.D. provided strategic advice and revised the manuscript. MHubert set up spatial analysis pipelines. E. B. and N. Gi provided transcriptomic and clinical data for the CURIMMUNE cohort. VP supervised some bioinformatics analysis. JV collected samples as the Principal Scientific Investigator of the LUNG PREDICT_LATE cohort, designed experiments, provided funding, supervised the research and wrote the manuscript.

## Acknowledgements

We would like to thank the staff of the core facilities and the BRC (Biological Resources Center) at the Cancer Research Center of Lyon (CRCL). Elisa Gobbini was supported by ESMO (any views, opinions, findings, conclusions or recommendations expressed in this material are those solely of the authors and do not necessarily reflect those of ESMO) and ITMO INSERM. We would like also to thank our financial supports : INSERM, INCA-DGOS PRTK_2017-022, INCA PLBIO INCa_4508, ANRS, ARC sign’it 2019, Ligue contre le Cancer (Labelisation, Régionale Auvergne-Rhône-Alpes et Saône-et-Loire, Comité de la Savoie, Comité de l’Ain), the Région Auvergne-Rhône-Alpes, SIRIC project (LYRIC, grant no. INCa_4664) and the LABEX DEVweCAN (ANR-10-LABX-0061) of the University of Lyon, within the program “Investissements d’Avenir” organized by the French National Research Agency (ANR) and the BMS foundation. This study has been partially supported through the grant EUR CARe N°ANR-18-EURE-0003 in the framework of the Programme des Investissements d’Avenir and an Eiffel Excellence doctoral fellowship to M. Hurtado.

## Data Availability Statement

The data generated in this study are available upon request from the corresponding author. The code to reproduce the analysis and figures is available on github at https://github.com/VeraPancaldiLab/LungPredict_DC_paper.

---

**Supplementary Table 1**: Samples features

**Supplementary Table 2** is provided as an .xlsx file containing 10 sheets with the output files produced in this study.

- **TF_matrix:** Matrix of TF activity scores inferred using VIPER and curated transcriptional regulatory networks (CollecTRI) for all samples.
- **TFs_modules:** TF modules inferred with their representative scores across samples.
- **Pathways activity:** Pathway activity scores inferred using the PROGENy database across samples.
- **Chemokines_scores:** Chemokine activity scores calculated for each sample based on predefined chemokine gene sets.
- **DC_signatures_scores:** DC signatures scores calculated for each sample based on predefined gene sets.
- **ARACNE_network:** Context-specific transcriptional regulatory network inferred using ARACNe-AP in the discovery cohort, reported as TF–target gene interactions with associated mutual information scores and p-values.
- **TF_matrix_ARACNE:** Matrix of TF activity scores inferred using VIPER based on the ARACNe-derived regulatory network.
- **GSVA_TF_signatures:** GSVA enrichment scores for TF signatures.
- **TF_matrix_CURIEIMMUNO:** Matrix of TF activity in the CURIEIMMUNO cohort, scores inferred using VIPER based on TF regulons curated using ARACNE.
- **GSVA_TF_signatures_CURIEIMMUNO:** GSVA enrichment scores for TF signatures in the CURIEIMMUNO cohort.

**Supplementary Figure 1.**
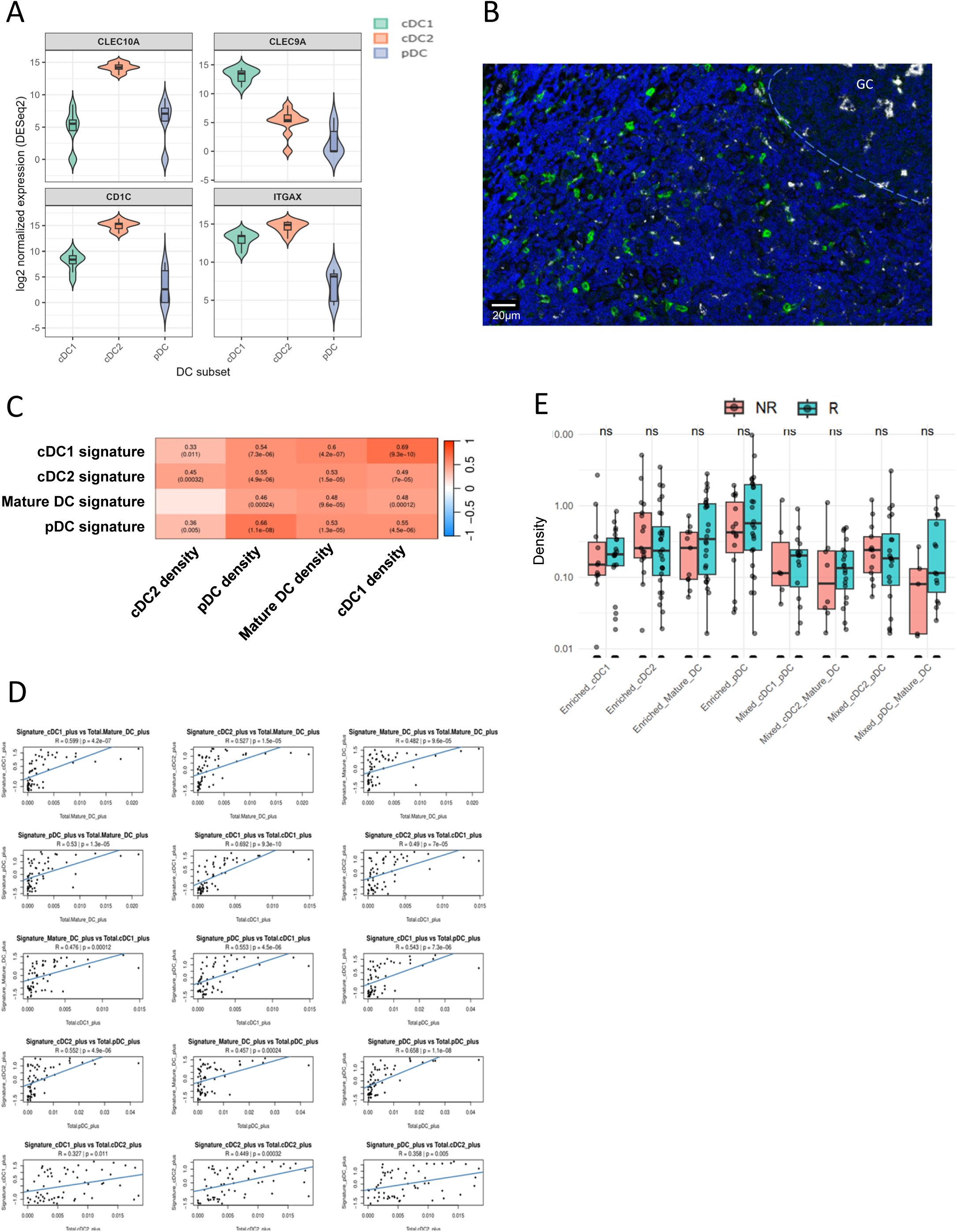
**(A)** log2 normalized expression of markers used for DC identification from tumor isolated DC microbulk RNAseq. **(B)** Representative multiplex immunofluorescence image of a lung cancer tissue stained for CD68 (white) and CLEC10A (green). Double-positive cells (macrophages) displayed lower CLEC10A intensity compared to CLEC10A single-positive cells (cDC2). Based on this, the annotation tool was calibrated to select the higher CLEC10A-expressing cells as true cDC2. **(C)** Heatmap showing Pearson correlation coefficients between DC subset densities defined by immunofluorescence and corresponding DC-specific transcriptional signatures by RNA-sequencing in the study cohort (N=60). The color scale represents the R values. **(D)** Scaterplot showing DC subset density (obtained *in situ* by mIF) with DC signature score obtained from RNAseq and linear regression**. (E)** Boxplot comparing DC megacluster densities between responder (n= 38) and non-responder patients (n=30). Statistical significance was assessed using the Wilcoxon test. ns = not significant.

**Supplementary Figure 2.**
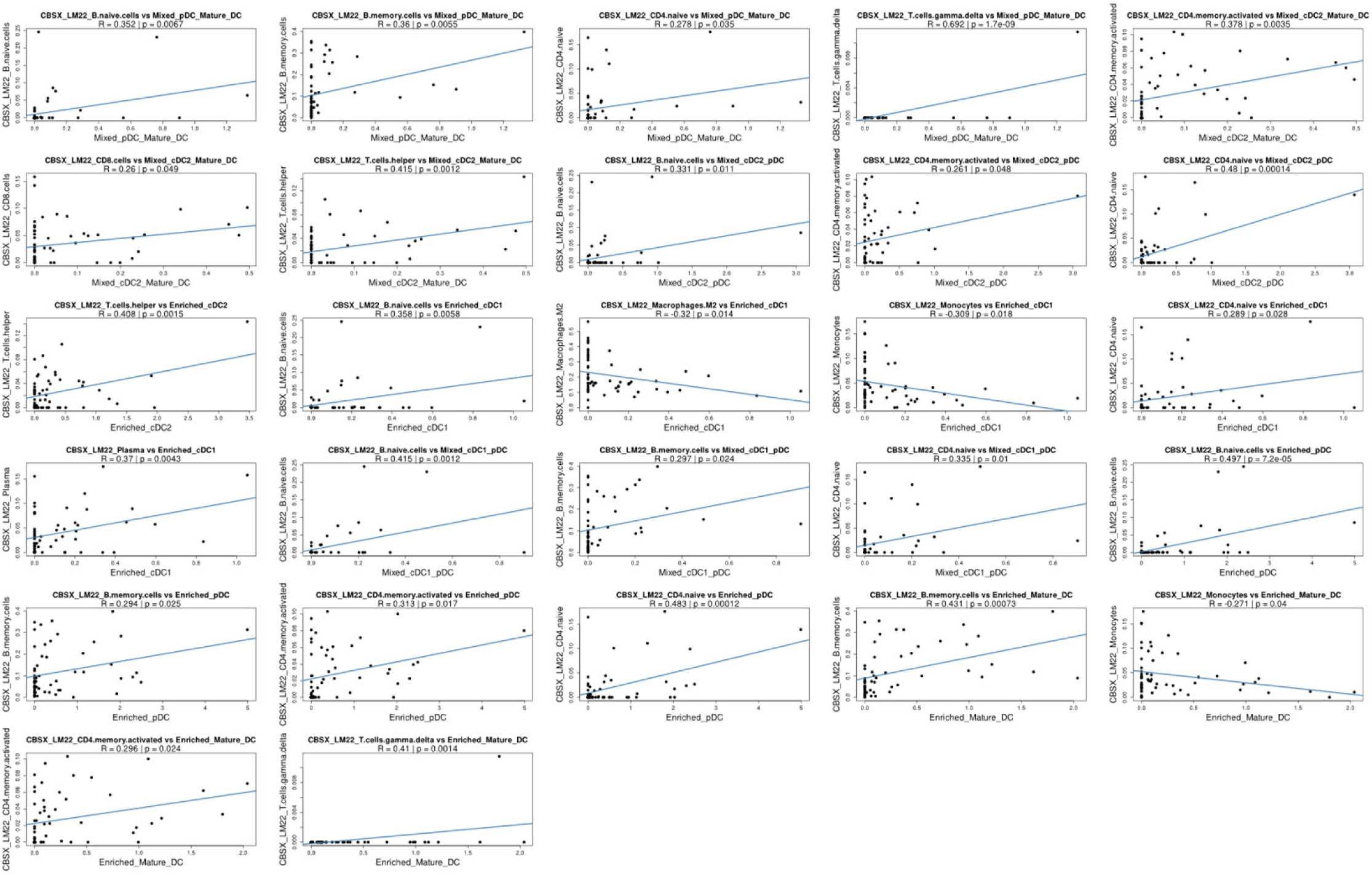
Scatter plot showing déconvolution scores obtained by CIBERSOTx (Y axis) with megacluster density (X axis).

**Supplementary Figure 3:**
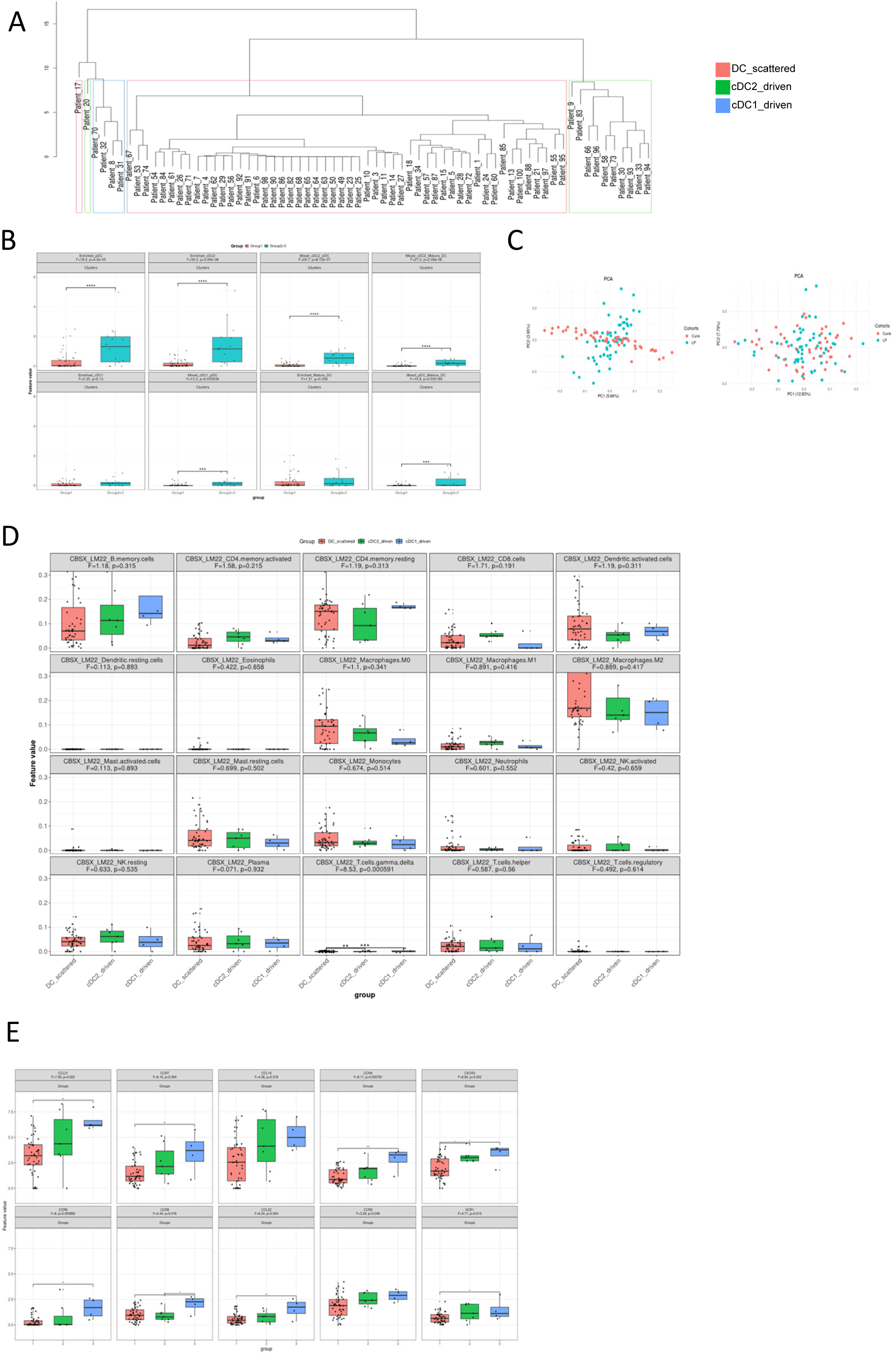
**(A)** Dendrogram of hierarchical clustering (ward.D2) (N = 68) based on the density of DC-derived megaclusters. Three major patient groups were identified: DC-scattered, cDC2-driven, and cDC1-driven. Two additional clusters were excluded from the analysis as they each included only one patientTFs differentially enriched in the cDC2-driven group compared to the others. **(B)** Boxplot showing the number of megacluster in group 1 (DC scaterred) versus cDC1 and cDC2 driven patients (group2+3). **(C)** Principal Component Analysis (PCA) performed on RNA-sequencing data from the study cohort (LUNG PREDICT) and the validation cohort (CURIMMUNE). PCA plots are shown for the entire transcriptome (left) and for TFs only (right). While PCA based on the whole transcriptome revealed a pronounced batch effect, this effect was absent when restricting the analysis to TFs.. **(D)** Boxplots showing the expression of immune cell population signatures (CIBERSORTx_LM22) across spatial groups. **(E)** Boxplots depicting chemokine and chemokine receptor expression across spatial groups. For all boxplots, statistical significance was assessed using one-way ANOVA, followed by Tukey’s honest significant difference (HSD) post hoc test for multiple comparisons: p ≤ 0.05 (*), p ≤ 0.01 (**), p ≤ 0.001 (***), p ≤ 0.0001 (****).

## Notes

### Competing Interest Statement

EG: speaker bureau for Merck Sharp & Dohme, Bristol-Myers Squibb, Regeneron, Sanofi, Janssen. Spouse of an AstraZeneca employee. Other authors declare no conflicts of interest.

